# Prenatal environment is associated with the pace of cortical network development over the first three years of life

**DOI:** 10.1101/2023.08.18.552639

**Authors:** Ursula A. Tooley, Aidan Latham, Jeanette K. Kenley, Dimitrios Alexopoulos, Tara Smyser, Barbara B. Warner, Joshua S. Shimony, Jeffrey J. Neil, Joan L. Luby, Deanna M. Barch, Cynthia E. Rogers, Chris D. Smyser

**Author notes:** Correspondence: Ursula A. Tooley.

## Abstract

Environmental influences on brain structure and function during early development have been well-characterized. In pre-registered analyses, we test the theory that socioeconomic status (SES) is associated with differences in trajectories of intrinsic brain network development from birth to three years (*n* = 261). Prenatal SES is associated with developmental increases in cortical network segregation, with neonates and toddlers from lower-SES backgrounds showing a steeper increase in cortical network segregation with age, consistent with accelerated network development. Associations between SES and cortical network segregation occur at the local scale and conform to a sensorimotor-association hierarchy of cortical organization. SES-associated differences in cortical network segregation are associated with language abilities at two years, such that lower segregation is associated with improved language abilities. These results yield key insight into the timing and directionality of associations between the early environment and trajectories of cortical development.

## Introduction

The first years of life are a time of rapid brain development, with intrinsic cortical organization becoming refined and large-scale brain systems developing into their adultlike configuration^1,2^. As children mature, intrinsic cortical networks become more segregated^3,4^, with sets of brain regions displaying more densely interconnected patterns of connectivity and large-scale systems becoming increasingly distinct^2,5–7^. This refinement of cortical organization occurs in a spatiotemporally patterned manner, with maturation occurring earlier in primary sensory regions than transmodal association regions^8^.The development of cortical network segregation also has implications for cognition, with higher levels of cortical network segregation associated with better cognitive abilities in adolescents and adults^6,9–12^. One environmental factor that is associated with the development of cortical network segregation — as well as many later life outcomes, including physical wellbeing^13,14^, cognitive ability^15^, and mental health^16^ — is socioeconomic status (SES).

Effects of SES on the development of cortical network segregation have been predominantly examined in older children and adolescents. In youth ages 8-22 years, SES moderated age-associated increases in cortical network segregation, such that youth from higher-SES backgrounds started with lower cortical network segregation but showed a steeper increase in cortical network segregation during adolescence than youth from lower-SES backgrounds, thus ending with higher levels of cortical network segregation at 22 years^17^.

Effects of SES on cortical network segregation were also found in youth ages 6-17 years living in low-SES neighborhoods^18^. Collectively, these studies suggest that low SES may accelerate the pace of cortical network segregation earlier in development, setting the stage for associations observed in older children and adolescents^19^. More broadly, associations between SES and intrinsic brain organization are visible as early as the first month of life^20,21^, but as of yet no longitudinal studies have examined how SES influences the development of cortical network segregation during the critical first years of life.

The pace of early brain development has been associated with important risk factors and behavioral outcomes. Changes in the pace of brain development have been linked to psychiatric disorders^22–24^, and accelerated pubertal development is associated with poorer mental health during adolescence^25–27^. An accelerated pace might also result in earlier declines in brain plasticity, curtailing the development of cortical circuitry optimally suited to the environment^28,29^. We have specifically posited that effects of SES on the pace of structural brain development might be reflected in the trajectory of functional brain development: more protracted structural brain development observed in children from higher-SES backgrounds may give rise to a longer, slower trajectory of intrinsic cortical network segregation^19^. If so, children from high-SES backgrounds should show more widespread connectivity and lower cortical network segregation early in development before the rapid development of a more segregated network architecture in later childhood and adolescence, leading to more effective cortical networks in adulthood^19^. Models drawn from evolutionary developmental frameworks also suggest that the early environment might affect the pace of maturation. These models posit that early experiences tailor organisms to anticipated conditions in the future and suggest that harsh and unpredictable environments may lead to life-history strategies resulting in accelerated development, while safe and nurturing environments may result in prolonged developmental strategies^30–32^. Relatedly, the recent “change of tempo” model contends that in situations of deprivation, such as inadequate nutrition or parental care, delaying maturation lowers children’s physiological requirements, while in the context of threat or abuse accelerated development may boost children’s ability to provide for immediate unmet safety needs^33^. At the same time, a recent review concluded that SES may simply be associated with alterations in brain development trajectories, not specific to pace, though this review noted the dearth of studies on intrinsic brain network development^34^. However, limited empirical work has employed the longitudinal data necessary to model effects of SES on trajectories of cortical network segregation, particularly in the earliest stages of life.

In this work, we explicitly test whether early SES is associated with differences in the pace of intrinsic cortical network segregation during the first three years of life. We capitalize on a unique cohort of neonates and toddlers with longitudinal neuroimaging data and extensively characterized early environments^35^. In this cohort, prenatal SES is associated with brain structure at birth^36,37^, however, associations between prenatal SES and brain function or longitudinal brain development have not yet been examined. Here, we test the theory that prenatal SES is associated with differences in the pace of functional brain network development, and that neonates and toddlers from higher-SES backgrounds might show a more protracted trajectory of cortical network segregation. In a set of pre-registered analyses, we examine the development of cortical network segregation during the first three years of life, and the moderating effects of prenatal SES on trajectories of cortical network segregation. We take a hierarchical approach to the question, first examining measures of cortical network segregation at the whole brain resolution, then analyzing at the level of functional brain systems, and finally at the level of individual brain regions. Next, we explore whether environmental effects are cortical-hierarchy-dependent based on the hypothesis that age-dependent changes in cortical plasticity might result in regional variability in associations between early SES and cortical network segregation. Finally, we examine whether differences in measures of cortical network segregation at two years of age are associated with language and cognitive abilities at the same age.

## Results

### Cortical network segregation increases from birth to three years

We first investigated the maturation of cortical functional network architecture between birth and three years of age by examining measures of cortical network segregation. Cortical network segregation can be measured at different scales, ranging from the local, or regional, scale to the global, or whole-brain, scale (***Figure 1a-c***). Global segregation captures the extent to which sets of systems in a cortical network are distinctly partitioned, while meso-scale segregation captures the extent to which a network can be divided into distinct subnetworks, and local segregation quantifies clustered connectivity at the regional or parcel level. We fit generalized additive mixed models (GAMMs) to measures of network segregation at each of these three levels, controlling for sex, amount of uncensored data included, in-scanner motion, and average network connectivity, and including a smooth term for age, where the smooth function (model fit for age) describes the relationship between cortical network segregation and age. Across scales, cortical network segregation increases with age. Global segregation, calculated from system segregation, increased with age (***Figure 1***, *F_s(age)_* = 6.26, *p* = 0.001, *p_FDR_* = 0.001), as did meso-scale segregation (*F_s(age)_* = 3.93, *p* = 0.01, *p_FDR_* = 0.01), and local segregation (*F_s(age)_* = 43.23, *p <* 0.0001, *p_FDR_* < 0.0001), indicating increasing refinement of network architecture during early development.

**Figure 1.**
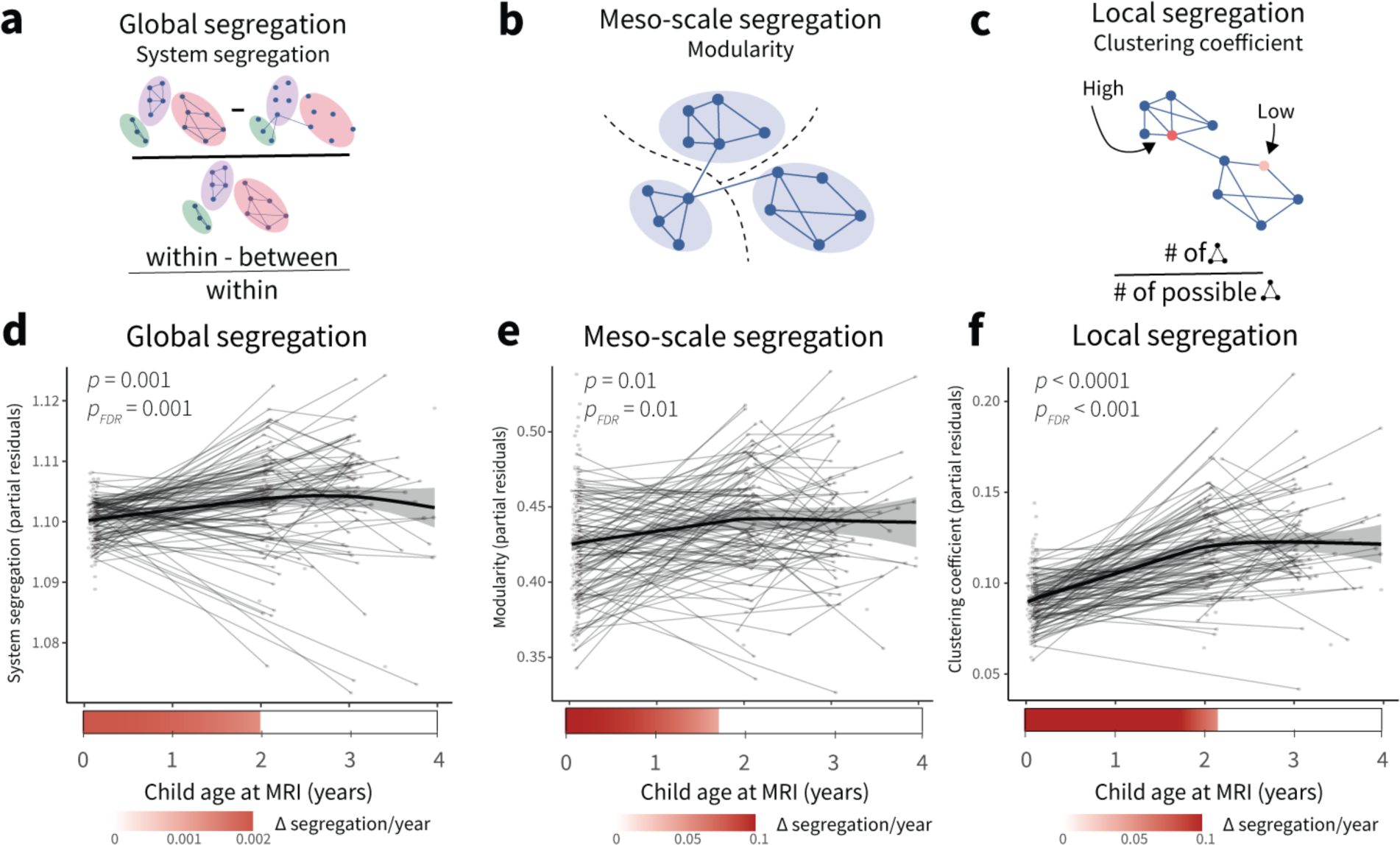
Cortical network segregation increases with age during the first three years of life. **a,** System segregation is a whole-brain measure of functional network segregation that quantifies the difference between mean within-system connectivity and mean between-system connectivity as a proportion of mean within-system connectivity. **b,** Modularity is a measure of network segregation that estimates the extent to which the nodes of a network, or in this case brain regions, can be subdivided into modules characterized by strong, dense intramodular connectivity and weak, sparse intermodular connectivity. Note that the modules are data driven, not *a priori* defined as functional systems. **c,** The clustering coefficient is a measure of local segregation that quantifies the amount of connectivity between a node and its neighbors. A node has a high clustering coefficient when a high proportion of its neighbors are also strongly connected to one another. In a weighted network, the clustering coefficient measures the strength of triangles around a node. **d,** Global segregation increases significantly with age. **e,** Meso-scale segregation increases significantly with age. **f,** Local segregation increases significantly with age. Individual points represent individual scans, with lines indicating scans from the same participant. Bars below the x-axis depict the derivative of the fitted smooth function of age. The filled portion of the bar indicates periods where the magnitude of the derivative of the fitted curve is significant, with the saturation of the fill representing the value of the derivative. Panels **a-c** reprinted with permission from ^7^.

Next, we probed the variation in the rate of change of cortical network segregation during the first three years of development. Here, the first derivative of the smooth function of age represents the rate of change in network segregation at a given developmental time point. Analysis of the derivatives of the age trajectories showed that measures of network segregation are increasing most strongly during the first two years of life. Global segregation increased between 0-1.93 years, meso-scale segregation increased between 0-1.70 years, and local segregation increased between 0-2.13 years, consistent with the most rapid change in cortical network segregation occurring early in development.

### Prenatal SES moderates trajectories of cortical network segregation

As described above, theoretical models posit that environmental influences on brain development might arise by way of effects on the pace of brain development, such that brain development proceeds faster in neonates and toddlers from lower-SES backgrounds. To test this hypothesis, we examined associations between the early environment and developmental increases in cortical network segregation. We used GAMMs to formally model age-by-SES interactions, which estimate how relationships between prenatal SES, our measure of early environment, and network segregation vary continuously with age. We observed significant and similar patterns of interactions between prenatal SES and age across multiple scales, such that infants and toddlers from lower-SES backgrounds show a faster increase in cortical network segregation than infants and toddlers from higher-SES backgrounds, ending up at a higher level of network segregation. This pattern held true for global segregation ***(Figure 2a***, *F_s(agexSES)_* = 6.38, *p* = 0.001, *p_FDR_* = 0.003), meso-scale segregation (***Figure 2b,*** *F_s(agexSES)_* = 9.86, *p* < 0.0001, *p_FDR_* = 0.0001), and local segregation (***Figure 2c,*** *F_s(agexSES)_* = 13.40, *p* < 0.0001, *p_FDR_* < 0.0001).

**Figure 2.**
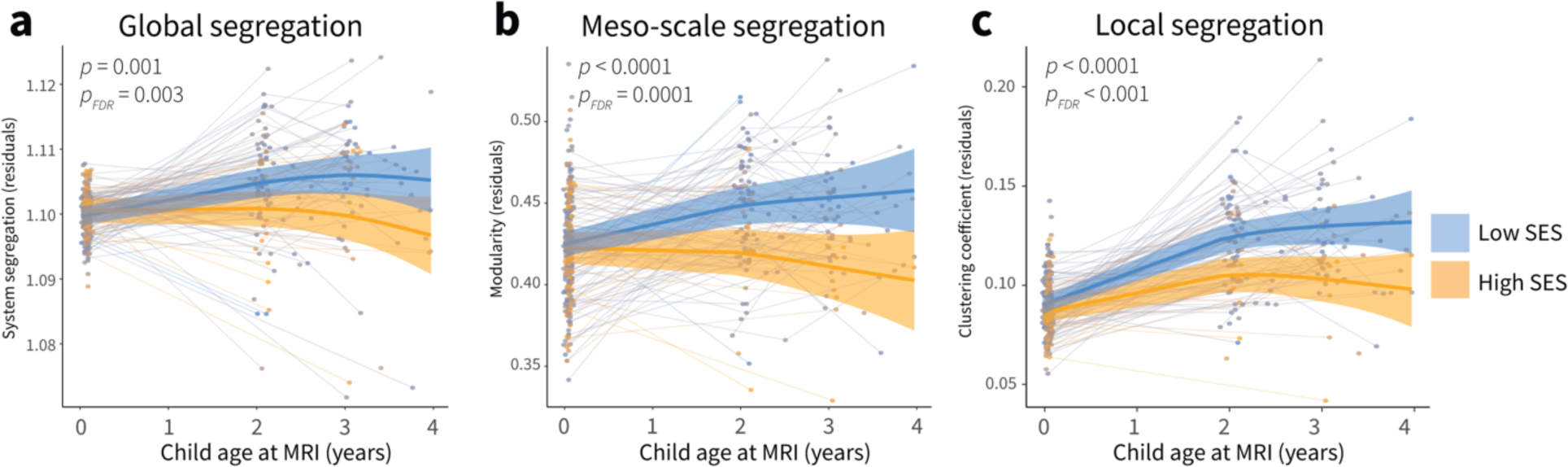
Associations between the early environment and developmental increases in cortical network segregation. **a,** Prenatal SES moderates trajectories of global cortical network segregation. **b,** Prenatal SES moderates trajectories of meso-scale cortical network segregation. **c,** Prenatal SES moderates trajectories of local cortical network segregation. Plots display fitted network segregation trajectories from GAMM models plotted by age for participants from high SES backgrounds (orange) and low SES backgrounds (blue). SES was modeled continuously, for visualization purposes here we show model trajectories from lowest and highest deciles of SES. Individual points represent individual scans, with lines indicating scans from the same participant.

Intuitively, measures of network segregation at different scales will be related: segregation at the global scale will encompass segregation at both the meso (intermediate) scale and the local scale. Correspondingly, we observe that measures of segregation are correlated with each other (|*r*|’s = 0.33-0.70, *p*’s < 1 × 10^-15^). To determine which effect drives the observed age-by-SES effects on cortical network segregation, we conducted a series of analyses examining age-by-SES effects on network segregation at each scale, controlling for network segregation at other scales. When we included average local segregation in the model for global segregation, age x SES no longer significantly predicts global segregation, suggesting that the SES-associated variance in age trajectories is contained in our measure of local segregation (*p* > 0.05). The same is true when we included average local segregation in our model for meso-scale segregation: the age-by-SES effect is no longer significant (*p* > 0.05). The inverse is not the case — including meso-scale or global segregation in our model of local segregation does not affect the significance of the age-by-SES interaction (*p*’s < 0.01).

Therefore, we concluded that the fundamental driver of the age-by-SES effects is variation in local network topology, as indexed by our measure of local segregation. Thus, in the analyses that follow we focus specifically on local segregation and associated variation in local network architecture.

### SES effects are strongest in somatomotor and dorsal attention systems

To characterize environmental effects on cortical network development in individual cortical regions, we conducted a series of exploratory (not pre-registered) analyses. We fit region-specific GAMM models to regional measures of local segregation, including a smooth term for age, and allowing age to interact with prenatal SES. A total of 56% of regions showed significant age-by-SES effects on local segregation, indicating that moderating effects of the environment on cortical network development are widespread across the cortex (*p_FDR_* < 0.05, ***Figure 3a).*** To provide insight into the magnitude of moderating effects of SES across cortical regions, we calculated the magnitude of variance explained by the addition of the age-by-SES interaction (*F*-statistic). The magnitude of SES effects on developmental increases in local segregation differed across the cortex, signifying that there is variability in the effect of the early environment on cortical network maturation across the developing cortex.

**Figure 3.**
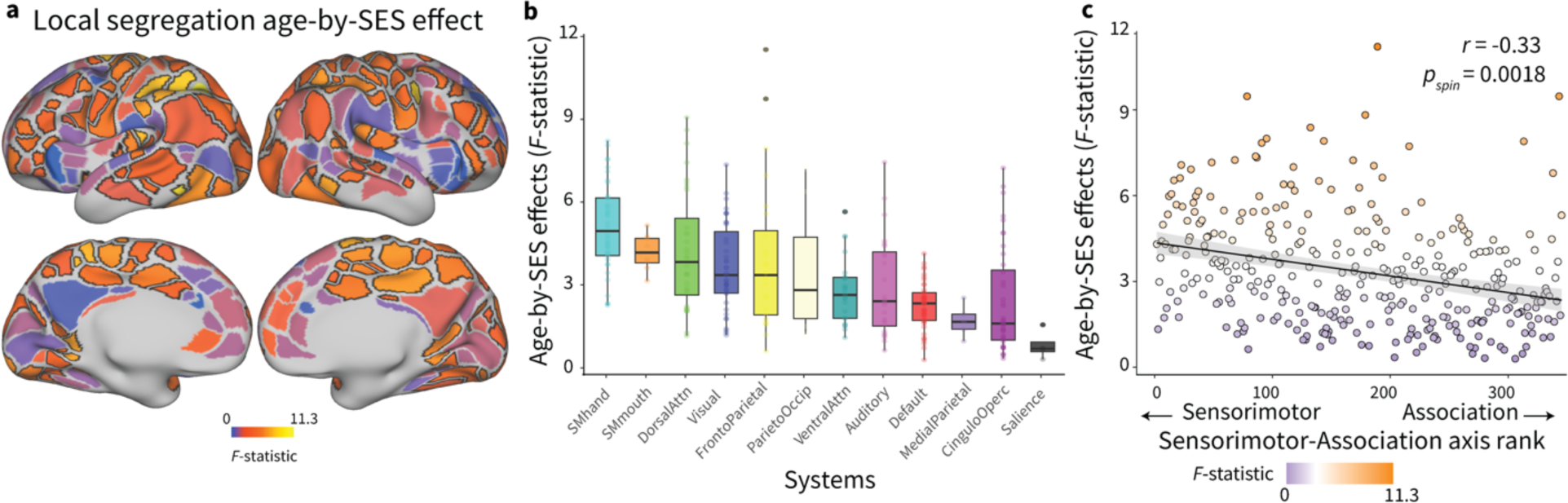
Environmental effects on developmental increases in local segregation are enriched in somatomotor systems. **a,** The heterogenous patterning of the magnitude of age-by-SES effects (*F-*statistic) on local segregation is shown on the cortical surface. Regions that show significant age-by-SES effects passing FDR correction at *p_FDR_* < 0.05 are outlined in black. **b,** SES effects on developmental increases in local segregation are enriched in somatomotor systems. Boxplots show median and interquartile range of the magnitude of age-by-SES effects; each point is an individual parcel. **c,** The magnitude of age-by-SES effects is related to a canonical sensorimotor-to-association axis of cortical organization. Each point is an individual parcel, the color of the points represents the magnitude of age-by-SES effects. A Spearman’s correlation between these two measures assessed by a conservative spin-based rotation test was significant (*r* = -0.33, *p_spin_* = 0.0018). The negative linear relationship between these two measures is plotted with a 95% confidence interval, colors of the points correspond to the magnitude of age-by-SES effects on regional local segregation.

To better understand the pattern of variability in environmental effects across the cortex, we asked whether moderating effects of prenatal SES on age-related increases in local segregation might be more pronounced in specific functional systems. The magnitude of SES effects on developmental increases in local segregation differed across functional systems (*H(12)* = 90.413, *p* < 1 × 10^-13^), with the strongest effects found in somatomotor-hand, somatomotor-mouth, dorsal attention, visual, and frontoparietal systems (***Figure 3b)***, consistent with environmental influences on cortical network development being most pronounced in early-developing sensorimotor regions during the first years of life. We wondered whether this was an overall principle of associations between the early environment and cortical network development, so we examined the correspondence between SES effects on developmental increases in local segregation and a canonical axis of sensorimotor-association cortical organization^8^. Age-by-SES effects on local segregation were negatively correlated with sensorimotor-association axis ranks across regions (***Figure 3c,*** *r* = -0.33, *p_spin_* = 0.0018), with larger environmental effects on developmental increases in local segregation characterizing the S-A axis’s sensorimotor pole, and smaller, near zero effects at the association pole.

### Cognitive consequences of SES-associated differences in local segregation

Finally, we sought to understand whether environmental influences on the development of cortical network segregation relate to differences in language and cognition that we observe later in development. SES-associated differences in language (***Fig 4a***, *F_s(agexSES)_* = 7.54, *p* < 0.0001, *p_FDR_* < 0.0001) and cognitive *(F_s(agexSES)_* = 12.65, *p* < 0.0001, *p_FDR_* < 0.0001) composite scores are evident from two years of age. Thus, we examined whether differences in local segregation at two years, when we first observe diverging trajectories of cortical network segregation associated with SES, were associated with language or cognition composite scores at the same time point. Average whole-cortex local segregation was not associated with cognition scores (*β* = -0.03, *p* = 0.766, *p_FDR_* = 0.766), but was negatively associated with language scores (***Figure 4b,*** *β* = -0.23, *p* = 0.017, *p_FDR_* = 0.033), indicating that lower levels of cortical network segregation seen in higher-SES toddlers are associated with better language abilities at two years of age. Importantly, when examining year two language ability and additionally controlling for prenatal SES, we find that both prenatal SES (*β* = -0.23, *p* = 0.027) and average local segregation (*β* = -0.25, *p* = 0.017) are significantly associated with language scores, indicating that cortical network segregation is independently associated with language ability at two years, above and beyond the association between early SES and later language abilities.

**Figure 4.**
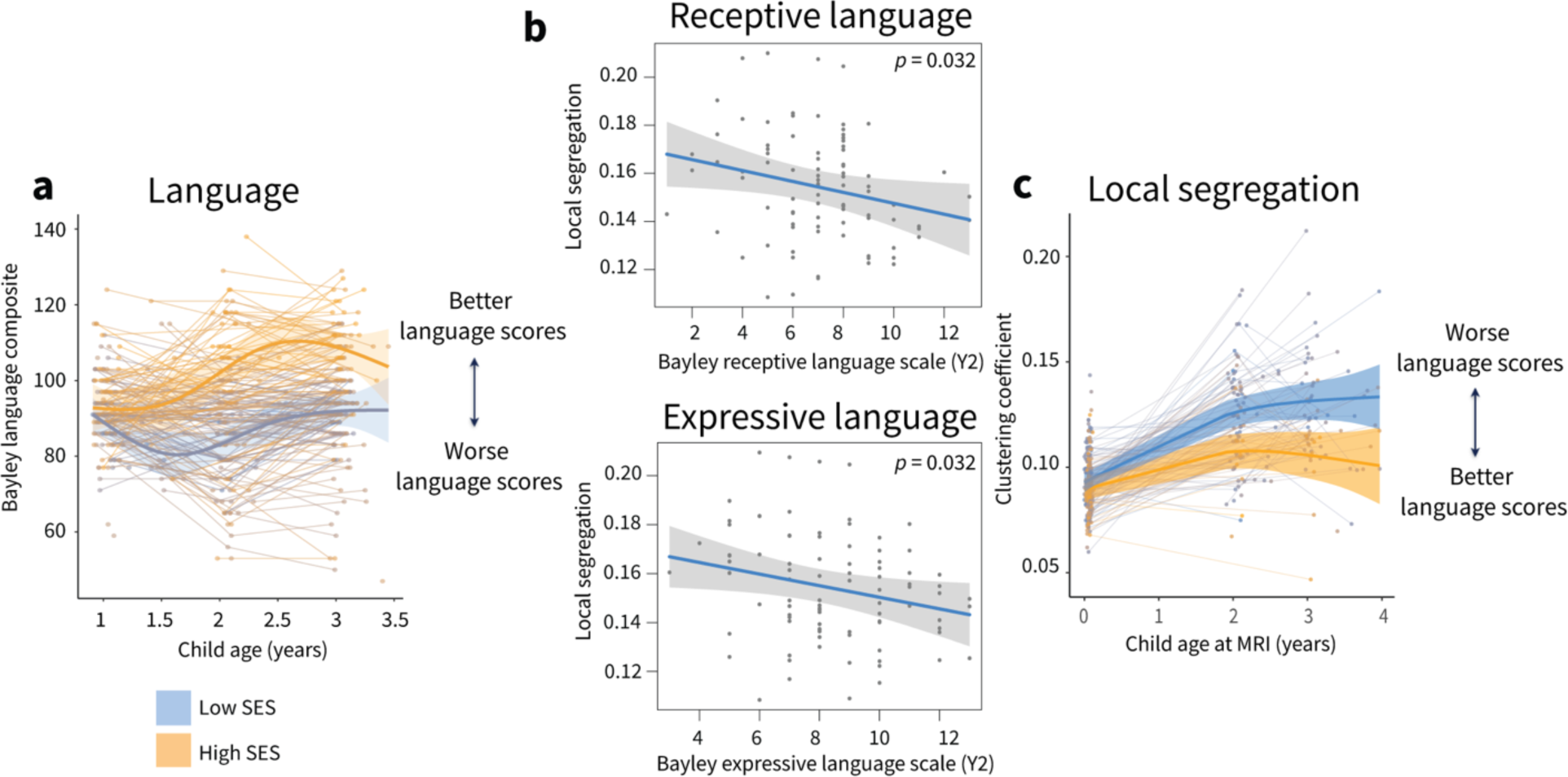
Associations between local cortical network segregation and language abilities at age two years. **a,** Effects of prenatal SES on trajectories of age-standardized Bayley language composite scores. SES was modeled continuously, for visualization purposes here we show model trajectories from lowest and highest deciles of SES. Individual points represent scores from individual participants, with lines indicating data collected from the same participant. **b,** Local segregation at two years of age was negatively associated with scores on both expressive and receptive language subscales. Scores on the receptive and expressive language subscales had different ranges, with children scoring slightly higher on the expressive language scale (x-axis). **c,** The directionality of this association is such that higher levels of local segregation, which are found in toddlers from lower-SES backgrounds, are associated with worse performance on measures of language ability.

We examined whether associations between local segregation and language abilities were driven by the receptive or expressive language subscales of the language composite, and found that local segregation was negatively associated with both expressive (*β* = -0.21, *p* = 0.032) and receptive language abilities (*β* = -0.21, *p* = 0.032). When controlling for prenatal SES, local segregation was associated with receptive language (*β* = -0.25, *p* = 0.022), but not expressive language (*β* = -0.21, *p* = 0.053), suggesting a domain-specific association of local segregation with receptive language abilities independent of the effect of SES at two years of age (***Fig 4c***).

While our sample size was reduced at the three-year timepoint due to COVID-related limitations in data collection (*n* = 90 at year two, *n* = 66 at year three), we additionally examined whether differences in local segregation at age three years were associated with language or cognition composite scores. We found a similar pattern of effect sizes at the three-year timepoint, with average local segregation negatively associated with language scores (*β* = -0.33, *p* = 0.006, *p_FDR_* = 0.01), but not cognitive scores (*β* = -0.20, *p* = 0.10, *p_FDR_* = 0.10). Local segregation was negatively associated with both receptive (*β* = -0.29, *p* = 0.015) and expressive language abilities (*β* = -0.35, *p* = 0.005). When controlling for prenatal SES at the three-year timepoint, we find similar effect sizes for associations between local segregation and language scores as at age 2, but neither associations with the language composite (*β* = -0.19, *p* = 0.14), or the expressive (*β* = -0.17, *p* = 0.17) or receptive (*β* = -0.19, *p* = 0.14) language scales are significant (***Fig 4c***).

### SES effects on developmental increases in cortical network segregation are robust to methodological variation

To ensure that the environmental effects on changes in cortical network segregation observed were robust to methodological variation and potential confounds, we performed five sensitivity analyses. We first conducted one of our pre-registered follow-up analyses, and evaluated whether effects of prenatal SES on developmental increases in cortical network segregation were accounted for by an alternate measure of the early environment. In this analysis, GAMMs were rerun with a composite measure of income-to-needs ratio (INR) and maternal education at birth rather than the disadvantage factor score (i.e., excluding neighborhood disadvantage, diet, and insurance status). When using this composite measure of SES, global segregation (*F_s(agexSES)_* = 3.74, *p* = 0.024, *p_FDR_* = 0.03), meso-scale segregation (*F_s(agexSES)_* = 7.78, *p* = 0.0005, *p_FDR_* = 0.0009), and local segregation (***Figure 5a,*** *F_s(agexSES)_* = 11.61, *p* < 0.0001, *p_FDR_* < 0.0001) all show significant and similar patterns of interactions, such that infants and toddlers from lower-SES backgrounds show a faster increase in cortical network segregation than infants and toddlers from higher-SES backgrounds. The magnitude of SES effects on developmental increases in local segregation continued to vary across the cortex (***Figure 5b***), with the strongest effects found in somatomotor-hand, visual, dorsal attention, and somatomotor-mouth systems (***Figure 5c***).

**Figure 5.**
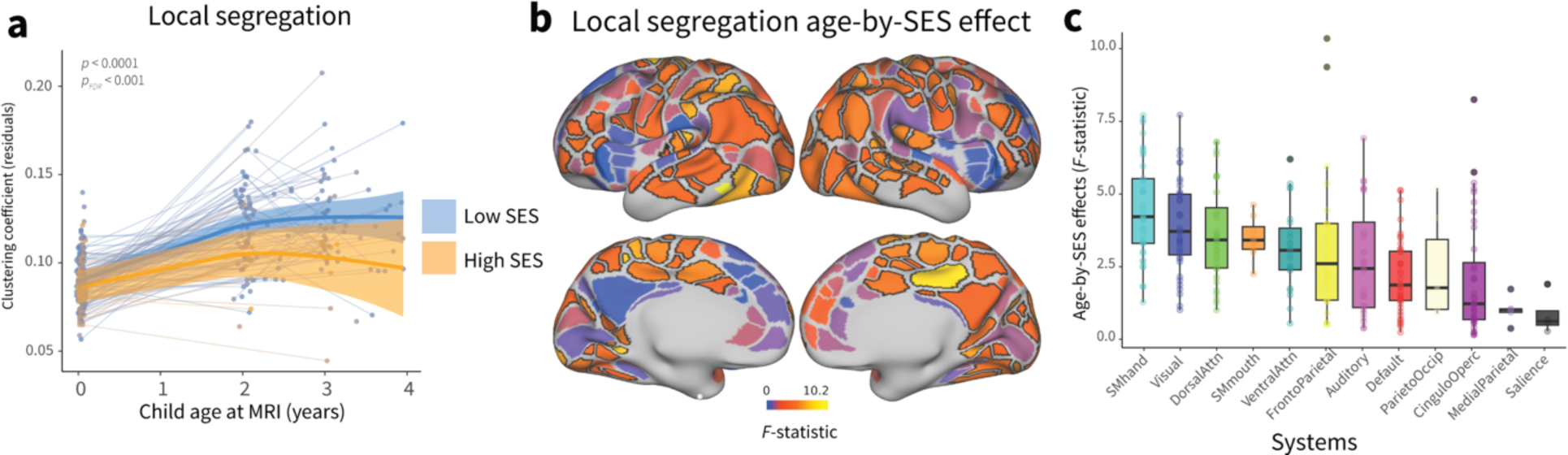
Associations between a composite measure of SES (maternal education and income-to-needs ratio) and developmental increases in cortical network segregation. **a,** Prenatal SES moderates trajectories of local cortical network segregation. **b**, The heterogenous patterning of the magnitude of age-by-SES effects (*F-*statistic) on local segregation is shown on the cortical surface. Regions that show significant age-by-SES effects passing FDR correction at *p_FDR_* < 0.05 are outlined in black. **c,** SES effects on developmental increases in local segregation are enriched in sensorimotor systems. Boxplots show median and interquartile range of the magnitude of age-by-SES effects; each point is an individual parcel.

We also evaluated whether effects of prenatal SES on developmental increases in cortical network segregation were accounted for by differences in sample composition over the study period, alterations in functional network architecture associated with head motion, or changes in SES over the study period. In each sensitivity analysis, environmental effects on developmental increases in cortical network segregation closely mirrored those observed in the main analysis, with neonates and toddlers from lower SES backgrounds showing a steeper increase in cortical functional network segregation than those from higher SES backgrounds, and environmental effects on developmental increases in cortical network segregation enriched in sensorimotor systems (see ***Supplementary Results & Figures 1-3***). These analyses verify that findings concerning the nature and patterning of environmental effects on developmental changes in cortical functional network segregation are robust to methodological variation.

### No evidence for environmental effects on age-associated changes in network integration

The previous analyses focused on developmental trajectories of network segregation, but another property of cortical network architecture that changes markedly as children develop is cortical work integration. Cortical network integration refers to the extent to which information can be integrated across multiple brain systems, and high levels of both network integration and network segregation together constitute the unique property of small-worldness found in adult brain networks^38^. Thus, we also pre-registered examining developmental changes in and age-by-SES effects on cortical network integration, as assessed by the average whole-brain participation coefficient. The participation coefficient is a measure of network integration that quantifies the diversity of the connections of a node across systems^39^, and has been linked in older children and adolescents to developmental changes in network segregation^7,40,41^ (***Figure 6a***).

**Figure 6.**
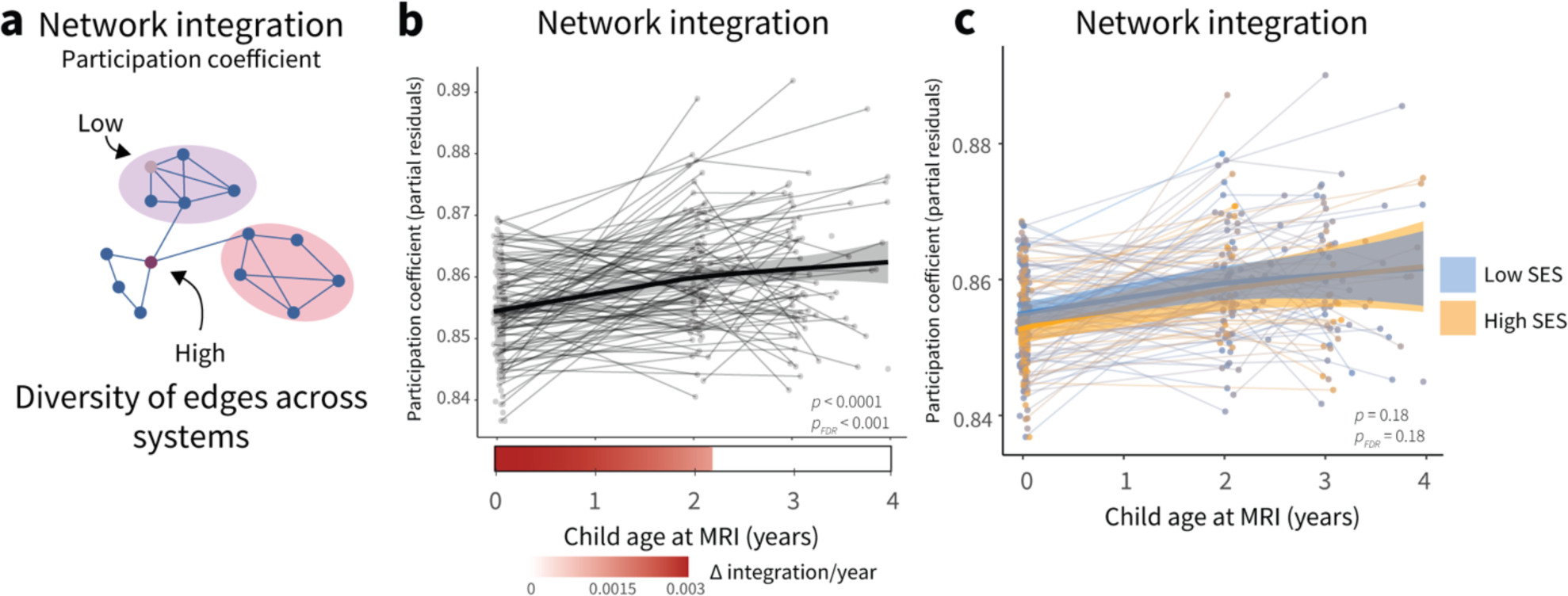
No evidence for environmental effects on age-associated changes in network integration. **a,** The participation coefficient is a measure of network integration that quantifies the diversity of the connections of a node. A node has a high participation coefficient when it is evenly connected to many different systems. Reprinted with permission from ^7^. **b,** Network integration increases significantly with age. Individual points represent individual scans, with lines indicating scans from the same participant. Bars below the x-axis depict the derivative of the fitted smooth function of age. The filled portion of the bar indicates periods where the magnitude of the derivative is significant, with the saturation of the fill representing the value of the derivative. **c**, Prenatal SES does not significantly moderate age-related increases in network integration.

As previously, we fit a GAMM to average whole-cortex network integration with a smooth term for age, where the smooth function (model fit for age) describes the relationship between network integration and age. Average network integration increased with age (***Figure 6b***, *F_s(age)_* = 14.11, *p* < 0.0001, *p_FDR_* < 0.0001), with the period of significant change occurring from birth to 2.27 years, consistent with the most rapid change occurring early in development. We found no evidence for moderating effects of prenatal SES on developmental changes in network integration (***Figure 6c,*** *F_s(agexSES)_* = 1.71, *p* = 0.18, *p_FDR_* = 0.18) in these primary analyses nor in supplemental analyses examining the alternative composite measure of SES described above (*F_s(agexSES)_* = 1.53, *p* = 0.12, *p_FDR_* = 0.12, see ***Supplemental Information***).

## Discussion

The first years of life are a critical window during which the development of the cortex proceeds in a heterochronous manner: sensory and motor cortices mature earlier than association cortices, and hierarchical refinement of plasticity-regulating features produces declines in the malleability of the cortex to environmental input, suggesting that the early environment may have a disproportionate effect on the pace of cortical development. In the current work, we show that the development of cortical brain networks during the first three years of life is strongly associated with features of the early environment, suggesting these influences may play a key role in shaping this trajectory. Specifically, we observe that developmental increases in cortical functional network segregation are accelerated in neonates and toddlers from disadvantaged backgrounds. Environmental effects on the development of cortical network segregation conform to a sensorimotor-association hierarchy of cortical organization, with the strongest effects visible on early-developing somatosensory systems that have high plasticity during this period. Differences in cortical functional network segregation associated with the early environment are also associated with language abilities at age two years when controlling for early SES.

We find that being born into a more advantaged (higher SES) environment is associated with a more protracted trajectory of cortical functional network development in toddlerhood and early childhood. Prior work has found that differences in intrinsic functional connectivity associated with SES are evident at 6 months of age^21^. Lower levels of cortical network segregation in children from more advantaged backgrounds at three years of age are also consistent with our earlier work in adolescents, where children from more advantaged backgrounds had lower levels of cortical network segregation at age 8 years^17^. This underscores the possibility that being born into a higher SES household may be associated with more protracted cortical functional network development throughout childhood, with children from high SES backgrounds showing more widespread connectivity and thus lower segregation early in development before the rapid development of a more segregated network architecture occurring during adolescence and into adulthood^19^. Experiences associated with low SES may result in earlier declines in plasticity in a heterogenous and spatially patterned manner across the cortex. For example, one study using a putative measure of plasticity to probe how developmental variation in intrinsic cortical activity is associated with the environment finds that lower SES is associated with reduced intrinsic activity, suggestive of decreased plasticity, in association cortex during adolescence^8^. Earlier reductions in plasticity associated with SES might thereby result in decreased synaptic proliferation^42,43^ or alterations in the development of inhibitory interneurons^44,45^, as has been found in animal models of early environmental stress, resulting in decreased range for optimal synaptic pruning and wiring of segregated cortical networks. In our cohort, neonates from more advantaged backgrounds show increased mean diffusivity, a measure of white matter microstructure, at birth compared to neonates from less advantaged backgrounds^46^. As this measure of white matter microstructure typically decreases with age^47^, we speculate that this pattern could be indicative of more protracted white matter development in neonates from more advantaged backgrounds; accelerated development associated with low SES might be observable in both intrinsic cortical network segregation and also the white matter connectivity that supports intrinsic brain activity, questions that can be explored in future work.

We find that alterations in cortical network segregation associated with SES are driven fundamentally by effects at the local scale. Associations between cortical network segregation and SES in older children and adolescents are also driven by alterations in local network architecture^17^. In youth and adolescents, effects of household SES on local segregation are evident in the prefrontal cortex specifically among youth living in low-SES neighborhoods, more evidence for environmental influences at the local scale during development^18^. However, adulthood SES affects aging-associated declines in global segregation^48,49^, thus one possibility is that effects of SES on cortical network segregation follow a progression from local to global across the lifespan. However, the authors did not examine local or meso-scale measures of segregation, so this possibility has not yet been directly addressed; more work is needed to clarify the scale of environmental effects on measures of cortical network segregation.

We find that environmental effects on the development of cortical functional network segregation conform to a sensorimotor-association hierarchy of cortical organization, with the strongest effects of the early environment visible in early-developing somatosensory systems, specifically, somatomotor, dorsal attention, and visual systems, with the frontoparietal system also showing strong environmental effects. While the dorsal attention system may not canonically be considered an early-developing system, dorsal attention regions mature dramatically between birth and two years of age^5^, are structurally and functionally connected to visual cortex^50,51^, and fall at the unimodal end of the primary gradient of macroscale cortical organization^52^. Environmental effects on cortical network segregation aligning with the sensorimotor-association hierarchy is consistent with a model where age-dependent changes in cortical plasticity allow environment effects to exert differential inputs on the human brain, depending on developmental timing, such that early plasticity in primary sensory regions allows for the largest environmental influences on intrinsic cortical activity in these regions during the first years of life.

Accumulating evidence from large-scale studies of older children and adolescents also suggests another possibility: environmental effects may be transduced through lower-level sensory systems, such that regardless of developmental stage, the strongest effects of the environment on intrinsic cortical activity are visible in somatosensory systems^53^. Across multiple indicators of SES, strong associations between SES and measures of functional connectivity are consistently found in somatomotor systems in children ages 9-11 years of age^54–57^. In youth and adolescents ages 8-21 years, SES associations with age-associated increases in cortical network segregation are also evident in somatomotor cortex^17^. It is also possible that due to the low level of social mobility in the United States population from which these studies were drawn^58,59^, SES later in development measured in these studies is nearly identical to participant’s SES at birth, and thus the effects measured are simply long-lasting instantiations of the effects of the birth environment on early-developing functional brain systems. Alternatively, brain areas at the sensorimotor pole of the S-A axis show the lowest variability between individuals^60,61^, and thus may enable the detection of effects above and beyond interindividual variability. While environmental associations were most prominent in the somatomotor system, we also observe associations between the early environment and cortical network development in the frontoparietal system. Interestingly, the environmental associations observed in the frontoparietal system are driven by strong age-by-SES interaction effects in regions in the intraparietal sulcus and middle temporal gyrus, rather than in lateral prefrontal regions, suggestive of alignment with an anterior-to-posterior developmental axis.

We find that SES-associated differences in cortical functional network segregation are also associated with language abilities at age two years, suggesting that environmental influences on the development of cortical network segregation might underlie SES-associated differences in language abilities observed later in development. The association between cortical network segregation and language abilities holds even when controlling for prenatal SES, suggesting that cortical network segregation is associated with language ability independent of the well-documented association between early SES and later language ability. While to our knowledge no other studies have specifically examined associations between cortical network segregation and language during early development, one study of toddlers and children ages 0-6 years found that increased between-system connectivity was associated with better language performance^62^, and infants with more language interactions had lower within-system connectivity in the language system^63^. Both of these findings potentially indicate that better language performance and more language exposure may be associated with lower levels of network segregation. The directionality of associations between cortical network segregation and language ability we find in toddlers is broadly reversed compared with associations found in adolescents and adults, where higher levels of cortical network segregation have been found to be advantageous for language and cognition^9,11,12^. During late childhood, children from higher-SES backgrounds show lower levels of cortical network segregation, but by early adulthood, higher SES is associated with higher cortical network segregation^17^, suggesting that initially low cortical network segregation might be beneficial for the development of cognitive and language abilities associated with higher SES^15^. However, the directionality of associations between cortical network segregation and behavior in adults also varies based on life stage and behavioral domain examined^6,10,64^, underscoring the importance of understanding developmental trajectories and domain specificity when interpreting the directionality of associations.

Several limitations and possible direct extensions of this work should be highlighted. First, it is important to acknowledge that socioeconomic disadvantage is but one form of disadvantage, and other forms not examined here, such as discrimination and systemic racism, may also be associated with brain development^65,66^. Different environmental exposures associated with SES — such as stress, cognitive stimulation, and unpredictability — also affect brain development differently^67–70^, and differentiating the relative contributions of these associated exposures to the effects that we observe is another critical and urgent future direction. Second, due to the impact of the COVID-19 pandemic, sample sizes at the toddler timepoints were smaller than originally intended, and thus our analyses of behavior at the two- and three-year time points should be considered exploratory; we share them as grounds for future work on the topic. Finally, while we observed associations between cortical intrinsic network development and language during toddlerhood, accelerated cortical network development may also have implications for the development of cognition and psychopathology that only become evident later in childhood. Longitudinal data from birth to middle childhood on both brain and behavioral development will be necessary to answer this question, data which will fortunately soon be available in the HEALthy Brain and Child Development (HBCD) study^71^.

The present study demonstrates that the development of intrinsic cortical brain networks during the first three years of life is associated with features of the early environment. The observed effects are consistent with an interpretation of accelerated cortical functional network development during the first years of life in neonates and toddlers from more disadvantaged backgrounds. Insight into the timing and directionality of environmental influences on trajectories of cortical brain development are crucial elements for understanding the optimal timing for interventions, to prevent cascading consequences of early maturation. Our results suggest that infancy and toddlerhood may be an important period for promoting healthy brain development, and emphasize the first years of life as a target for policies aimed at supporting optimal child development.

## Citation diversity statement

Recent work in several fields of science has identified a bias in citation practices such that papers from women and other minority scholars are undercited relative to the number of such papers in the field^72–78^. We used prior methods^76,79^ to measure that our references contain 18.75% woman(first)/woman(last), 22.47% man/woman, 30.28% woman/man, and 28.51% man/man authors. This method is limited in that a) names, pronouns, and social media profiles used to construct the databases may not, in every case, be indicative of gender identity and b) it cannot account for intersex, non-binary, or transgender people. We used additional methods^80,81^ to measure that our references contain 13.32% author of color (first)/author of color(last), 14.58% white author/author of color, 21.41% author of color/white author, and 50.69% white author/white author. This method is limited in that a) names, Census entries, and Wikipedia profiles used to predict racial/ethnic categories may not be indicative of racial/ethnic identity, and b) it cannot account for Indigenous and mixed-race authors, or those who may face differential biases due to the ambiguous racialization or ethnicization of their names. We look forward to future work that could help us to better understand how to support equitable practices in science.

## Acknowledgements

We thank the families involved with the study, and Karen Lukas, Eva Loveless, and Rich Nagel for assistance with scanning neonates. We thank the past and current team members of the March of Dimes Prematurity Research Center at Washington University in St Louis, the Early Emotional Development Program at Washington University in St Louis, and the Washington University Neonatal Development Research Group.

## Methods

### Participants

Neonates were recruited as a part of the Early Life Adversity, Biological Embedding, and Risk for Developmental Precursors of Mental Health Disorders (eLABE) cohort^35^, whose participants were recruited under the parent March of Dimes study^82^. Pregnant mothers were recruited and enrolled between the second and third trimesters. Recruitment oversampled mother-infant pairs facing adversity (e.g. poverty and stress).Inclusion criteria for the study included speaking English, mother age 18 years or older, and singleton birth (see ***Supplemental Table 1*** for detailed participant information). Women with alcohol or other substance abuse were excluded. Anatomic MR images were reviewed by a neuroradiologist (J.S.S.) and pediatric neurologist (C.D.S.). Subjects were excluded from the current analyses if they had evidence of brain injury or were born preterm (<37 weeks gestational age, GA).

Additional exclusion criteria included pregnancy complications (but not gestational diabetes or hypertension) and known fetal abnormalities including intrauterine growth restriction. This study was approved by the Human Studies Committees at Washington University in St. Louis and informed consent was obtained from a parent of all participants.

At each timepoint, all participants with usable functional magnetic resonance imaging (fMRI) and demographic data were included. At the neonatal timepoint, 385 neonates were scanned, and participants were excluded from all timepoints for the following reasons: <37 weeks GA at birth (*n* = 54), brain injury (*n* = 17), neonatal intensive care unit stay for > 7 days, required intubation or chest tube, antibiotics for > 3 days, cardiac disease or metabolic disorder (*n* = 36), birthweight < 2,000 g (*n* = 1), and IRB exclusion (*n* = 1). There were 306 participants who did not meet any of these exclusion criteria (note that some met multiple exclusion criteria). Of these participants, 261 participants (age range = 38-45 post-menstrual weeks, *M* = 41.3 months) were included in the current analyses at the neonatal timepoint, neonates were excluded for no usable T2 for registration (*n* = 27), no functional magnetic resonance imaging (fMRI) data collected or < 10 min of usable fMRI data after motion censoring (*n* = 12), or visible artifacts in FC data (*n* = 7).

At the two-year time point, 202 participants were scanned, of which 162 were healthy full-term neonates not subject to the exclusions above. Participants were additionally excluded for no usable T1 for registration (*n* = 68) or no functional magnetic resonance imaging (fMRI) data collected (*n* = 2), resulting in 92 participants (range = 1.91-2.61 years, *M* = 2.11 years) included at year two. At the three-year point, 132 participants were scanned, of which 98 were healthy full-term neonates. Participants were additionally excluded for no usable T1 for registration (*n* = 31) or < 5 min of usable fMRI data after motion censoring (*n* = 1) resulting in 66 participants included in the current analyses from year 3.

### MRI data acquisition

Imaging was performed without sedating medications at all three time points using a 3T Prisma scanner (Siemens Corp.) and 64-channel head coil. During the scan session, structural images were collected: a T2-weighted image at the neonatal timepoint (sagittal, 208 slices, 0.8-mm isotropic resolution, echo time, TE = 563 ms, repetition time, TR = 3200 ms) and a T1-weighted image at the two- and three-year timepoints (sagittal, 208 slices, 0.8-mm isotropic resolution, repetition time = 2400 ms, echo time = 2.22 ms). Resting-state functional imaging data (fMRI) were collected using a blood oxygen level–dependent (BOLD) gradient-recalled echo-planar multiband sequence (72 slices, 2.0-mm isotropic resolution, echo time = 37 ms, repetition time = 800 ms, multiband factor = 8, 420 volumes). Spin-echo field maps were obtained (at least 1 anterior–posterior and 1 posterior-anterior) during each session with the same parameters. Between 2 and 9 5.6 minute fMRI BOLD scans were acquired, depending on how the child tolerated the scan (range = 7.5 - 44.8 min). Framewise Integrated Real-time MRI Monitoring (FIRMM^83,84^) was used during scanning to monitor real time participant movement.

### MRI data preprocessing

fMRI preprocessing included correction of intensity differences attributable to interleaved acquisition, bias field correction, intensity normalization of each run to a whole-brain mode value of 1,000, linear realignment within and across runs to compensate for rigid body motion, and linear registration of BOLD images to the adult Talairach isotropic atlas performed in a single step. Neonates were registered: BOLD to individual T2 to group-average T2 from this cohort to 711-2N Talairach atlas. Toddlers were registered: BOLD to individual T1 to group-average T1 from this cohort to 711-2N Talairach atlas. Field distortion correction was performed, using the FSL TOPUP toolbox (http://fsl.fmrib.ox.ac.uk/fsl/fslwiki/TOPUP).

Following initial processing, the surface-based neonatal parcellation approach, Melbourne Children’s Regional Brain Atlases (MCRIB), was used to generate surfaces for each neonatal subject and the volumetric resting-state BOLD timeseries were mapped to subject-specific surfaces using established procedures adapted from the Human Connectome Project as implemented in Connectome Workbench 1.2.3. Freesurfer 7.2 was used to generate surfaces for each toddler subject, and the volumetric resting-state BOLD timeseries were mapped to subject-specific surfaces using established procedures adapted from the Human Connectome Project as implemented in Connectome Workbench 1.2.3.

After mapping to the surface, each dataset underwent resting-state fMRI preprocessing. In the initial iteration, the data were processed with the following steps: (i) demean and detrend within run, (ii) nuisance regression including white matter, ventricles, extra-axial cerebrospinal fluid, and whole brain, as well as 24-parameter Friston expansion regressors derived from head motion. Next, frames contaminated by motion were censored as described below. Finally, the initial rs–fc preprocessing stream was repeated on the output of the initial preprocessing using only the frames that had passed motion criteria, with the addition of interpolating censored frames and band-pass filtering (0.005 Hz < *f* < 0.1 Hz).

Neonatal fMRI data were censored at FD > 0.25 mm, with the additional restriction that only epochs of at least 3 consecutive frames FD < 0.25 mm were included. This FD threshold was selected after taking into account the smaller radius of infants’ heads^85^ and reviewing motion traces in several subjects^86,87^; respiratory filtering is unsuitable for neonatal fMRI data due to the higher respiratory rate of neonates. Toddler (two-year and three-year) fMRI data were censored based on a threshold of FD_filt_ > 0.2 mm, using a filtered framewise displacement trace corrected for the effect of respiration (FIRMM filtered FD^88,89^), with the additional restriction that only epochs of at least 3 consecutive frames FD < 0.2 mm were included.

In order to be included in the study, a minimum of 5 minutes (375 frames) of data retained after censoring was required, though 99% of scans across timepoints had > 10 min of data retained after censoring (*M* = 17.4 min (1308 frames), range = 7.1-41.9 min). To account for any potential patterns of FC related to head motion or amount of data included in analyses, we calculated (i) the number of frames retained after censoring and (ii) the average FD across uncensored frames for each individual subject and included these values as subject-level covariates in in all analyses. We additionally calculated (iii) the average FD across all frames for each individual subject, used as subject-level covariates in sensitivity analyses. Neither average uncensored FD nor total number of frames retained after censoring were related to prenatal disadvantage (uncensored FD: *r* = 0.014, *p* = 0.77; total number of frames: *r* = 0.06, *p* = 0.22).

fMRI data were aligned across subjects into the “fs_LR32k” surface space using spherical registration. Timecourses for surface data were smoothed with geodesic 2D Gaussian kernels (σ = 2.25 mm).

### Network analysis

Residual mean BOLD time series from each participant at each timepoint were extracted from a 333-region cortical parcellation^90^. The functional connectivity matrix was represented as a graph or network^91^. Network edges between nodes (brain regions) were represented by the Fisher *z*-transformed Pearson correlation between time courses from pairs of surface regions^92^. Because there is not yet consensus on the spatial layout of neonatal functional networks^85,93–98^, we assigned nodes to 13 large-scale functional systems (also sometimes called “networks”) based on the definitions of functional systems derived in healthy adults^90^.

Across the cortex, we calculated three summary measures of functional network segregation, which quantify the extent to which groups or subnetworks of nodes, or brain regions, display densely interconnected patterns of connectivity (see ***Figure 1***). System segregation, our measure of global segregation, quantifies the difference between mean within-system connectivity and mean between-system connectivity as a proportion of mean within-system connectivity^4,9^, given an *a priori* partition of nodes into systems, in this case the 13 large-scale functional systems^90^. Modularity, quantified by the modularity quality index (Q), is a measure of meso-scale network segregation that estimates the extent to which the nodes of a network can be subdivided into groups or modules characterized by strong, dense intramodular connectivity and weak, sparse intermodular connectivity. Local segregation was measured using the clustering coefficient, which quantifies the amount of connectivity between a node and its strongest neighbors^99–102^. A node has a high clustering coefficient when a high proportion of its neighbors are also strong neighbors of each other. Local segregation at the whole-brain level was calculated as the average of the clustering coefficient across all cortical regions. In later analyses of regional specificity, we examined the clustering coefficient of individual nodes or regions. We specifically chose measures of functional network architecture that were suitable for weighted, signed networks, when possible.

We also estimated a measure of functional network integration, the participation coefficient, which quantifies the diversity of a node’s connections across systems^39,103^. See ***Supplemental Methods*** for further details of network analysis and equations for measures of network segregation and integration.

### Demographics and socioeconomic status

Mothers completed surveys in each trimester, at delivery, and during follow up visits every 4 months to assess social background, mental health, and life experiences. As described in ^35^, prenatal SES was assessed using a latent factor of socioeconomic disadvantage from a confirmatory factor analysis that included measures of mother’s income-to-needs ratio, educational attainment, area deprivation index, insurance status, and nutrition. Maternal self-reported highest level of education and health insurance status were collected in trimester 1. Mothers reported household income and persons in the home to calculate INR in each trimester^104^. Home addresses were collected at delivery to obtain Area Deprivation Index percentiles; the area deprivation index is a geocoding measure that ranks neighborhoods by socioeconomic disadvantage compared with the national average based on census block data, including factors for the domains of income, education, employment, and housing quality^105^. Prior work in the eLABE cohort has shown that, while 26% of mothers changed addresses during pregnancy, there was no significant change in block groups during pregnancy^106^, and we similarly find that there was not significant social mobility (change in ADI) during the study time period in our sample (*F_s(child age)_* = 1.44, *p* = 0.19). Maternal nutrition was assessed in the third trimester or at delivery using the Healthy Eating Index, a validated dietary assessment tool available through the National Institutes of Health used to measure diet quality based on U.S. Dietary Guidelines for Americans^107^. Higher scores on the disadvantage factor are indicative of lower SES.

Data on demographic and socioeconomic indicators (education, household income, insurance status, ADI) were also collected during follow-up visits. Prenatal disadvantage was highly correlated with disadvantage at years 1-3 (*r*’s = 0.92-0.93, *p*’s < 1 × 10^-16^). While we focus our analyses on the disadvantage factor assessed at birth, supplementary analyses investigate the contribution of disadvantage at later time points. Additionally, we examine the effect of using a composite variable of parental education and income to assess SES, rather than the disadvantage factor, for consistency with prior literature, in our sensitivity analyses.

### Cognitive and language outcomes

The Bayley Scales of Infant and Toddler Development-Third Edition (Bayley-III) was used to assess cognitive, language, and motor abilities at both two and three years of age. Based on prior evidence that network segregation is associated with higher-order cognitive abilities in adults^9^, we specifically examined age-standardized cognitive and language composite scores (M = 100, SD = 15); the language composite consists of the Receptive Language and Expressive Language subscales. At the two-year timepoint, *n* = 90 children had usable Bayley assessments and fMRI data (2 children with imaging data were excluded for low-quality or missing Bayley). At the three-year timepoint, *n* = 66 children had usable Bayley assessments and fMRI data (no children with imaging data were excluded for low-quality or missing Bayley).

### Data analysis

Our analyses relating age and SES to cortical functional network architecture were pre-registered at https://aspredicted.org/eb5pd.pdf. Any deviations from the original plan or additional exploratory analyses have been fully described below.

### Statistical models

To flexibly model longitudinal linear and non-linear relationships between cortical functional network segregation and age, we implemented GAMMs (generalized additive mixed models) using the *mgcv* package in R^108^. GAMMs were fit with functional network segregation as the dependent variable, age as a smooth term, a random effect of participant, and biological sex assigned at birth, in-scanner motion (average framewise displacement), number of frames of fMRI retained after censoring, and average functional network weight (average network connectivity) as linear covariates. Each GAMM estimates a smooth function (the model age fit, generated from a linear combination of weighted basis functions) that describes the relationship between functional network segregation and age, thus modeling the developmental trajectory of network segregation. Four basis functions were specified as the maximum flexibility afforded to age splines in all models (*k* = 4). Models were fit using thin plate regression splines as the smooth term basis set and the restricted maximal likelihood approach for smoothing parameter selection. Average network weight was included to control for global differences in connectivity strength^109–111^. Random effects included a random intercept per participant. To test for windows of significant change across the age range, we calculated the first derivative of the smooth function of age from the GAMM model using finite differences, and then generated a simultaneous 95% confidence interval of the derivative^112^ using the *gratia* package in R. The first derivative of this smooth function represents the rate of change in network segregation at a given developmental time point. Intervals of significant age-related change were identified as areas where the simultaneous confidence interval of the derivative does not include zero. Multiple comparisons correction was applied across models of developmental changes in cortical functional network segregation using FDR correction^113^.

To examine moderating effects of SES on age-related increases in functional network segregation, we allowed the smoothed age effect in the GAMM to interact with SES; predictors thus included an age-by-SES interaction term, a smooth term for age, and covariates including sex, in-scanner motion, number of frames of fMRI, and average network weight. We compared two interaction models: a simpler varying coefficient (linear-nonlinear) model that allows the smooth term for age to vary as a linear function of SES, and a more complex non-linear interaction (bivariate smooth) model that allows the smooth term for age to vary as a fully non-linear function of SES. We compared models using Bayesian information criterion (BIC), and evaluated the significance of the interaction term for the selected model. All models were best fit using the simpler varying coefficient (linear-nonlinear) model that allows the linear association between SES and network segregation to vary as a smooth function of age. Interaction p-values were confirmed using a parametric bootstrap likelihood ratio test (*pbkrtest* package in R) for significance estimation in the mixed model context. Multiple comparisons correction was applied across models of age-by-SES effects on cortical functional network segregation using FDR correction.

To evaluate the scale at which SES affects measures of functional network segregation, we fit models for each of our measures of local, meso-scale, and global segregation in turn, controlling for each of the other measures of segregation. For example, we first fit a GAMM for the age-by-SES effect on system segregation while including modularity as a fixed effect, then fit a GAMM for age-by-SES effects on system segregation while including the clustering coefficient as a fixed effect.

To characterize regional specificity of SES effects on maturational changes in local segregation, we fit region-specific GAMMs with the same interaction structure and covariates as the whole-brain models. Models were fit separately for each parcellated cortical region. For each regional GAMM, the significance of the age-by-SES interaction term was assessed in a fixed degree-of-freedom context to ensure stable and accurate estimation. We corrected *p-* values across all region-wise GAMMs using FDR correction and set statistical significance at *p_FDR_* < 0.05. To establish the overall magnitude of the moderating effect of SES on age-associated increases in local segregation, which we refer to throughout as a region’s overall age-by-SES effect magnitude, we used the *F-*statistic for the age-by-SES interaction effect. To evaluate the age-by-SES effect across functional systems, we used the effect magnitude at each parcel within each system. A Kruskal-Wallis test was used to compare the magnitude of the age-by-SES effects across functional systems.

Moderating effects of SES on associations between age and language composite scores were also modeled with GAMMs; we allowed the smoothed age effect in the GAMM to interact with SES, using the same model comparison framework as above. We tested a varying-coefficient (linear-nonlinear) model and a more complex fully non-linear interaction (bivariate smooth) model, comparing models using Bayesian information criterion (BIC), and evaluated the significance of the interaction term for the selected model; the bivariate smooth model fit the language composite best, while the linear-nonlinear model fit the cognitive composite best. Note that as these the composite scores are standardized for age, the smoothed effect of age in this model solely accounts for variation over time in language scores relative to age norms.

When examining relationships between local cortical network segregation and language outcomes at the two-year and three-year timepoints, due to the smaller number of datapoints available, we used linear models (results are qualitatively unchanged when using GAMs). We estimated associations between local segregation and Bayley language scaled scores, controlling for biological sex assigned at birth, in-scanner motion (average framewise displacement), number of frames of fMRI retained after censoring, and average functional network weight. Multiple comparisons correction was applied across each set of two models using FDR correction.

### Comparison of cortical maps

A previously derived axis of sensorimotor-association cortical organization^8^ was retrieved from https://github.com/PennLINC/S-A_ArchetypalAxis. To quantify the association between S-A axis ranks and observed environmental effects on developmental increases in local segregation, we used Spearman’s rank correlations and tested for statistical significance using spin-based spatial permutation tests^114,115^, which account for spatial covariance structure common in neuroimaging data. We generated a null distribution based on 10,000 spherical rotations, and compared the observed value to the null.

### Code and data availability

All analyses were conducted in R4.1.2 (https://www.r-project.org/) and MATLAB R2021b. Functions from the Brain Connectivity Toolbox^103^ were used to calculate measures of network segregation and integration. Freely available MATLAB code from https://github.com/mychan24/system_matrix_tools was used to calculate system segregation. Surfaces and regional effects were shown on cortical surfaces generated by MCRIB using the *cifti* and *ciftiTools* packages^116^ and Connectome Workbench. Code for all analyses presented here is publicly available at https://github.com/utooley/Tooley2023_prenatal_env_cortical_network_dev. Deidentified data will be deposited into a public repository upon publication.

### Deviations from pre-registration

We did not conduct some analyses in the pre-registration that were deemed potential exploratory analyses and turned out to be unhelpful in elucidating the effects we found. Specifically, as the primary driver of the age-by-SES effect was found to be on local segregation (the clustering coefficient) and alterations in local topology, we did not examine age-by-SES effects on within- and between-system connectivity; this analysis would have been informative for probing age-by-SES effects on system segregation. As our measure of local segregation, the clustering coefficient, does not rely on an *a priori* assignment of regions to functional systems, we deemed an analysis of functional-system-level connectivity unhelpful in investigating the effects we found here. Also, in a follow-up analysis we had planned to examine the effect of including only participants whose disadvantage factor score did not change by > 1 SD between timepoints. However, as upon investigation there was little change in direct indicators of SES during the study period (see ***Supplementary*** Figure 3), and the composition of the disadvantage factor score changed from the birth to toddler timepoints (prenatal Healthy Eating Index was removed from the factor score), we decided not to pursue this analysis.

Additionally, we include here follow-up analyses that were not part of our pre-registration to probe the contributors and potential behavioral associations of the moderating effects of SES on cortical functional network development trajectories that we found in the pre-registered analyses. Specifically, we examined a) which scale of segregation was the fundamental driver of age-by-SES effects on cortical network segregation, b) regional variation in age-by-SES effects on local segregation, and c) associations between local segregation and language and cognitive outcomes. These analyses were not pre-registered and should be considered exploratory in nature.

Finally, we initially planned to rerun any GAMMs with estimated degrees of freedom (EDF) of the age smooth < 2 as linear mixed effects models, but upon visual inspection of developmental trajectories of cortical network segregation, non-linear models of age seemed appropriate even for those measures with age EDFs slightly under 2. As GAMMs can model both linear and non-linear relationships, in the case that there is no non-linear relationship between age and an outcome measure, the smooth term for age will be penalized down to a linear term. Thus, for ease of comparison and to avoid switching modeling approaches, we have instead used GAMMs for all models in the included analyses.

