## Supplemental Information for "Prenatal environment is associated with the pace of cortical network development over the first three years of life"

### Supplemental Methods

***MRI data acquisition.*** At the neonatal timepoint, after feeding, the infant was swaddled and positioned in a head-stabilizing vacuum fix wrap. A nurse familiar with neonate transport and resuscitation was present at all MRI scans. Heart rate and blood oxygenation were measured continuously throughout all scans, and infants were monitored visually via video. Based on visual monitoring through a camera, infants slept through scans as indicated by eye closure and minimal movements. Imaging was performed without sedating medications using a Siemens 3T Prisma scanner and 64-channel head coil. A T2-weighted image (sagittal, 208 slices, 0.8mm isotropic resolution, echo time, TE = 563 ms, repetition time, TR = 3,200 ms) was collected. For the resting-state fMRI, functional imaging was performed using a blood-oxygen-level dependent (BOLD) gradient-recalled echo-planar multiband (MB) sequence (72 slices, 2.0-mm isotropic resolution, TE = 37 ms, TR = 800 ms, MB factor = 8). Scans were collected in both the anterior-posterior (AP) and posterior–anterior (PA) direction; a typical session included 2 AP scans and 2 PA scans. Each scan was 420 frames, which is 5.6 min in length.

At the toddler (two- and three-year) timepoints, during naptime or after bedtime, the child was allowed to fall asleep naturally in the presence of the parent. The child was then positioned in the scanner with the use of custom head padding and weighted cushions and blankets. Toddlers were monitored visually via video; based on video, toddlers slept through scans as indicated by eye closure and minimal movements. Imaging was performed without sedating medications using a Siemens 3T Prisma scanner and 64-channel head coil. A T1-weighted image (sagittal, 208 slices, 0.8mm isotropic resolution, repetition time, TR = 2,400 ms, echo time, TE = 2.22 ms) was collected. For the resting-state fMRI, functional imaging was performed using an identical blood-oxygen-level dependent (BOLD) gradient-recalled echo-planar multiband (MB) sequence (72 slices, 2.0-mm isotropic resolution, TE = 37 ms, TR = 800 ms, MB factor = 8). Scans were collected in the anterior-posterior (AP) direction; the scans were 420 frames, which is 5.6 min in length.

***Network analysis.*** Residual mean BOLD time series were extracted from a 333-region cortical parcellation[1], and we represented the functional connectivity matrix as a graph or network[2]. Regions were represented by network nodes, and the functional connectivity between region *i* and region *j* was represented by the network edge between node *i* and node *j*. We estimated the functional connectivity between any two brain regions by calculating the product-moment correlation coefficient *r* between the mean activity time series of region *i* and the mean activity time series of region *j*[3]. We used this encoding of the data as a network to produce an undirected, signed, and weighted adjacency matrix A. Correlations were subsequently *r*-to-*z* transformed. Because there is not yet consensus on the spatial layout of neonatal functional networks[4–10], we assigned nodes to 13 large-scale systems (also sometimes called “networks”) based on the definitions of functional systems derived in healthy adults[1].

Prior evidence has demonstrated that the maintenance of edge weights is critical for an accurate understanding of the underlying biology of neural systems[11,12], and work in applied mathematics has demonstrated that graph-related calculations are markedly more robust in weighted graphs than in binary graphs[13]. In light of these two lines of evidence and recent work in the field developing methods sensitive to the topologies present in weak versus strong edges[14], we maintained all edge weights without thresholding and studied the full graph including both positive and negative correlations[15,16].

***Measures of functional network architecture***

*System segregation.* System segregation quantifies the difference in mean within-system connectivity and mean between-system connectivity as a proportion of within-system connectivity. Previous work has linked this measure to aging-related changes in brain networks and poorer cognitive ability across age[17,18]. In these analyses, we define system segregation as in [17], as:

$$\frac{\bar{a}_{within} - \bar{a}_{between}}{\bar{a}_{within}}$$

Where $\bar{a}_{within}$ is the mean edge weight between nodes within the same system and $\bar{a}_{between}$ is the mean edge weight between nodes of one system to all nodes in other systems. We assigned nodes to systems based on a 13-system partition[1]. Freely available MATLAB code from https://github.com/mychan24/system_matrix_tools was used to calculate system segregation.

*Modularity.* Statistics that quantify the modular structure of a network assess the extent to which a network's nodes can be subdivided into groups or modules characterized by strong, dense intramodular connectivity and weak, sparse intermodular connectivity. We considered the most commonly studied mesoscale organization—assortative community structure—that is commonly assessed by maximizing a modularity quality function[19,20]. Unlike system segregation, the modularity quality index is independent of a mapping of nodes to functional systems. Higher modularity is indicative of a more highly segregated network at the mesoscale. Our approach is built on the modularity quality function originally defined by [21] and subsequently extended to weighted and signed networks by various groups.

Specifically, we follow [14] by first letting the weight of a positive connection between nodes $i$ and $j$ be given by $a_{ij}^{+}$ , the weight of a negative connection between nodes $i$ and $j$ be given by $a^{-}{}_{ij}$ , and the strength of a node $i$, $s_i^{\pm} = \sum_j a^{\overset{\pm}{ij}}$, be given by the sum of the positive or negative $j$ connection weights of $i$. We denote the chance expected within-module connection weights as $e_{ij}^{+}$ for positive weights and $e_{ij}^{-}$ for negative weights, where $e_{ij}^{\pm} = \frac{s_i^{\pm} s_j^{\pm}}{v^{\pm}}$. We let the total weight $v^{\pm} = \sum_{ij} a_{ij}^{\pm}$ be the sum of all positive or negative connection weights in the network. Then, the asymmetric generalization of the modularity quality index is given by:

$$Q^* = \frac{1}{v^+}\sum_{ij}\left(a_{ij}^+ - e_{ij}^+\right)\delta_{M_iM_j} - \frac{1}{v^+ + v^-}\sum_{ij}\left(a_{ij}^- - e_{ij}^-\right)\delta_{M_iM_j}$$

where $M_i$ is the community to which node $i$ is assigned, and $M_j$ is the community to which node $j$ is assigned. We use a Louvain-like locally greedy algorithm as a heuristic to maximize this modularity quality index subject to a partition M of nodes into communities. We ran the Louvain algorithm 100 times per network, and detected on average four (*M* = 3.76, *SD* = 0.51) communities using modularity maximization in our developmental sample.

*Clustering coefficient***.** To assess local network segregation, we used a commonly studied graph measure of local connectivity—the clustering coefficient— that is commonly interpreted as reflecting the capacity of the system for processing within the immediate neighborhood of a given network node[22–25]. We specifically used a formulation that was recently generalized to signed weighted networks[26,27]. This version is sensitive to nonredundancy in path information based on edge sign as well as edge weight and importantly distinguishes between positive triangles and negative triangles, which have distinct meanings in networks constructed from correlation matrices.

We let the functional connectivity network of a single participant be represented as the graph $G = (V, E)$, where $V$ and $E$ are the vertex and edge sets, respectively. We let $a_{ij}$ be the weight associated with the edge $(i, j) \in V$, and define the weighted adjacency matrix of $G$ as $A = [a_{ij}]$. The clustering coefficient of node $i$ with neighbors $j$ and $q$ is given by:

$$C_i = \frac{\sum_{jq}\left(a_{ji}a_{iq}a_{jq}\right)}{\sum_{j\neq q}\left|a_{ji}a_{iq}\right|}$$

The clustering coefficient of the entire network was calculated as the average of the clustering coefficient across all nodes as follows:

$$C = \frac{1}{n}\sum_{i\in N} C_i$$

In this way, we obtained estimates of the regional and global clustering coefficient for each subject in the sample.

*Participation coefficient*. The participation coefficient is a measure of network integration that quantifies the diversity of a node's connections across communities, and has been linked in older children and adolescents to developmental changes in network segregation[28–30]. A node has a high participation coefficient when it is evenly and strongly connected to many different systems. In these analyses, we define the participation coefficient $P_i$ of a node $i$ as:

$$P_i = 1 - \sum_{k\in K}\left(\frac{a_{ik}}{s_i}\right)$$

where $k$ is a system in a set $K$ of systems, in this case defined by the *a priori* mapping of nodes to intrinsic functional systems, $a_{ik}$ is the positive (negative) weight of edges between node $i$ and nodes in system $k$, and $s_i$ is the positive (negative) strength of node *i*. The participation coefficient was calculated separately on negative and positive weights[31].

As in our analyses of local segregation, the participation coefficient of the entire network was calculated as the average positive (negative) participation coefficient across all nodes as follows:

$$P = \frac{1}{n}\sum_{i \in N} P_i$$

The average positive and negative participation coefficient for each participant's network were averaged to obtain a global measure of network integration.

### Supplemental Results

To ensure that the environmental effects on changes in cortical network segregation observed were robust to methodological variation and potential confounds, we performed five sensitivity analyses. The first sensitivity analysis is described in the main text. In the second sensitivity analysis, GAMMs were rerun with only participants who had full data at two or more timepoints (***Supplementary Figure 1***). In a third sensitivity analysis, we refit GAMMs including a measure of pre-censoring motion, rather than the post-censoring measure used in our main analyses (***Supplementary Figure 2***). Finally, we conducted several pre-registered follow-up analyses related to changes in SES over time. We first examined the extent of changes in indicators of SES, finding that there was little social mobility during the study period (***Supplementary Figure 3).*** Then, we reran GAMMs including the socioeconomic disadvantage factor calculated at Y1 and Y2, rather than at birth.

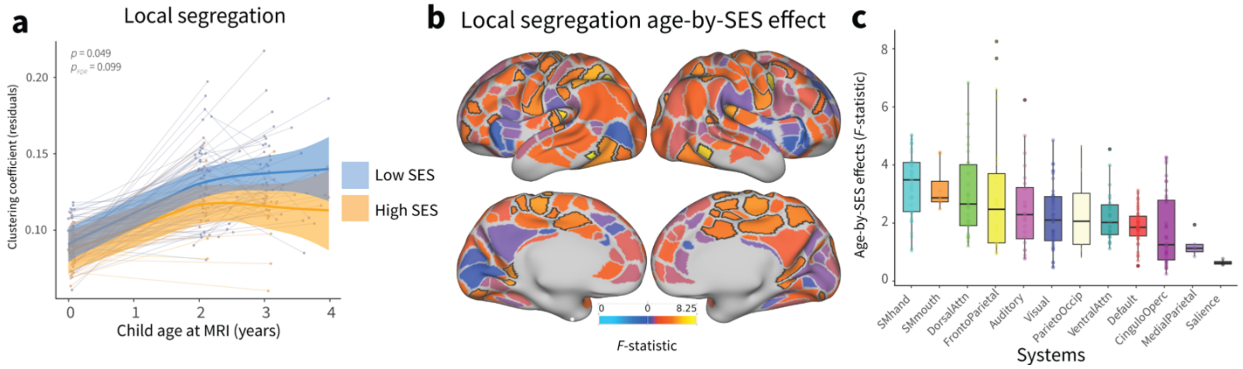


**Supplemental Figure 1**. Associations between the early environment and developmental increases in cortical network segregation, including only participants with two or more timepoints of data. **a,** Prenatal SES moderates trajectories of local cortical network segregation. **b,** The heterogenous patterning of the magnitude of age-by-SES effects (*F*-statistic) on local segregation is shown on the cortical surface. Regions that show significant age-by-SES effects passing FDR correction at $p_{FDR} < 0.05$ are outlined in black. **c,** SES effects on developmental increases in local segregation are enriched in sensorimotor systems. Boxplots show median and interquartile range of the magnitude of age-by-SES effects; each point is an individual parcel.

We additionally verified whether results are robust to only including participants with two or more timepoints of data collected. When limiting our sample to this subsample, due to a decrease in sample size (*n* = 132 unique participants), we have less power to detect significant effects, however, we still expect to observe effects in the same direction as those observed in our primary analyses. Global segregation ($F_{s(agexSES)}$ = 0.001, $p$ = 0.09, $p_{FDR}$ = 0.13), meso-scale segregation ($F_{s(agexSES)}$ = 9.24, $p$ = 0.0001, $p_{FDR}$ = 0.0005), and local segregation (***Supplemental Figure 1a,*** $F_{s(agexSES)}$ = 2.78, $p$ = 0.049, $p_{FDR}$ = 0.09) show similar patterns of interactions, such that infants and toddlers from lower-SES backgrounds show a faster increase in cortical network segregation than infants and toddlers from higher-SES backgrounds. We did not find evidence for moderating effects of SES on developmental changes in network integration ($F_{s(agexSES)}$ = 0.001, $p$ = 0.99, $p_{FDR}$ = 0.99). The magnitude of SES effects on developmental increases in local segregation differed across functional systems, with the strongest effects found in somatomotor-hand, somatomotor-mouth, dorsal attention, and frontoparietal systems (***Supplemental Figure 1c***).

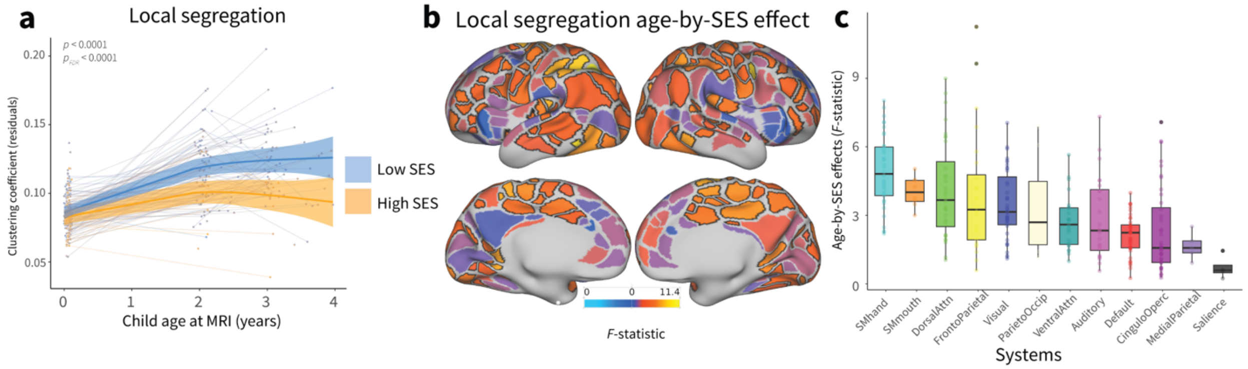


**Supplemental Figure 2.** Associations between the early environment and developmental increases in cortical network segregation, controlling for pre-censoring motion. **a,** Prenatal SES moderates trajectories of local cortical network segregation. **b,** The heterogenous patterning of the magnitude of age-by-SES effects (*F*-statistic) on local segregation is shown on the cortical surface. Regions that show significant age-by-SES effects passing FDR correction at $p_{FDR} <$ 0.05 are outlined in black. **c,** SES effects on developmental increases in local segregation are enriched in sensorimotor systems. Boxplots show median and interquartile range of the magnitude of age-by-SES effects; each point is an individual parcel.

We included a measure of pre-censoring motion, the total framewise displacement across all frames, instead of post-censoring motion, as a subject-level covariate. Global segregation ($F_{s(agexSES)}$ = 5.94, $p$ = 0.0029, $p_{FDR}$ = 0.0038), meso-scale segregation ($F_{s(agexSES)}$ = 9.70, $p <$ 0.0001, $p_{FDR}$ = 0.0002), and local segregation (***Supplemental Figure 2a,*** $F_{s(agexSES)}$ = 13.17, $p <$ 0.0001, $p_{FDR} <$ 0.0001) all show significant and similar patterns of interactions, such that infants and toddlers from lower-SES backgrounds show a faster increase in cortical network segregation than infants and toddlers from higher-SES backgrounds. We did not find evidence for moderating effects of SES on developmental changes in network integration ($F_{s(agexSES)}$ = 1.37, $p$ = 0.26, $p_{FDR}$ = 0.26). The magnitude of SES effects on developmental increases in local segregation differed across functional systems, with the strongest effects found in somatomotor-hand, somatomotor-mouth, dorsal attention, and frontoparietal systems (***Supplemental Figure 2c***).

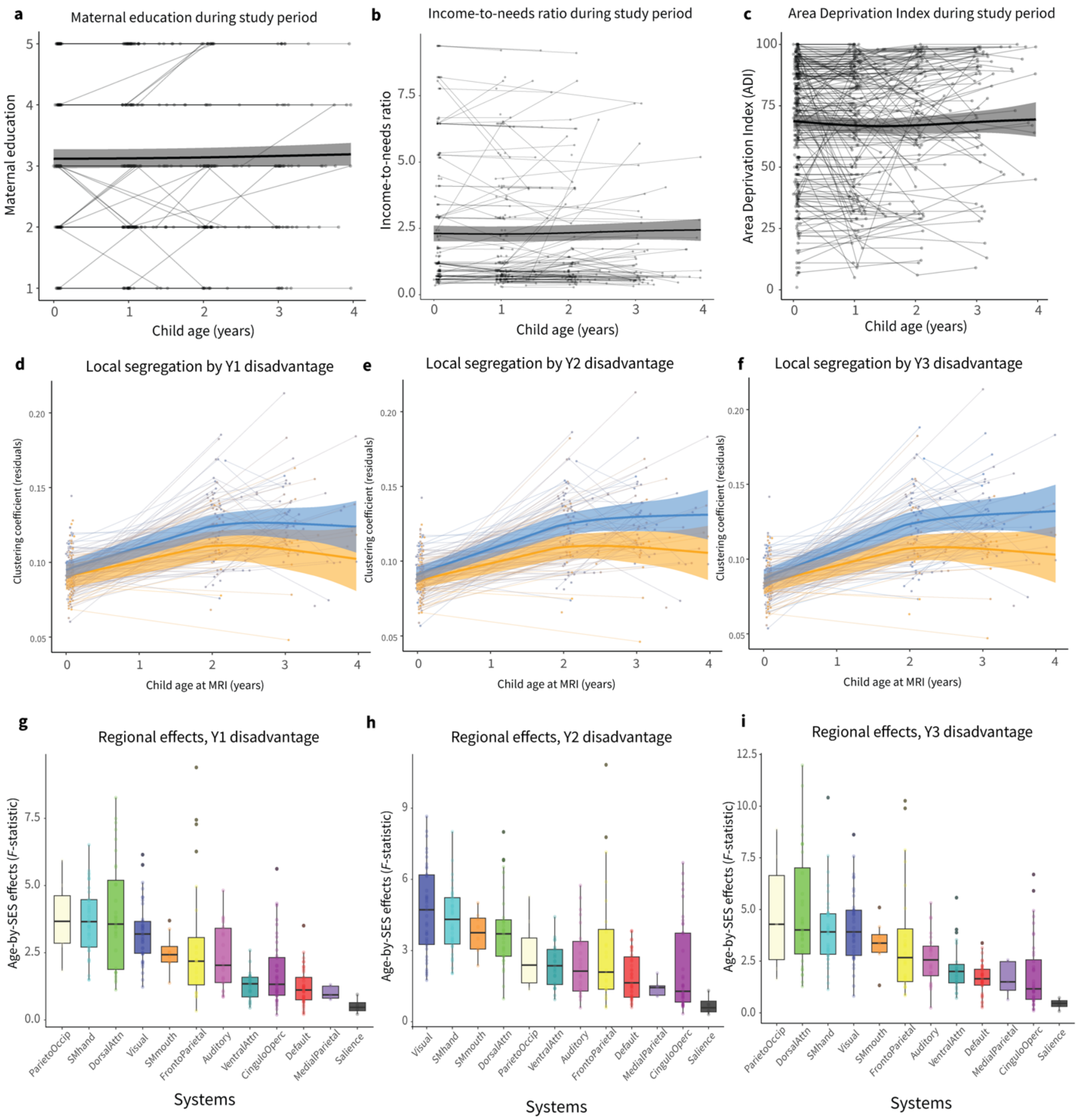


**Supplemental Figure 3.** Little evidence for changes in SES during the study period. **a,** Maternal education does not change significantly over the study period. **b,** Income-to-needs ratio does not change significantly over the study period. **c,** Area deprivation index (ADI) does not change significantly over the study period. **d,** Year one disadvantage moderates trajectories of local cortical network segregation. **e,** Year two disadvantage moderates trajectories of local cortical network segregation. **f,** Year three disadvantage moderates trajectories of local cortical network segregation. **g,** Year one disadvantage effects on developmental increases in local segregation across functional systems. **h,** Year two disadvantage effects on developmental increases in local segregation across functional systems. **i,** Year three disadvantage effects on developmental increases in local segregation across functional systems. Boxplots show median and interquartile range of the magnitude of age-by-SES effects; each point is an individual parcel.

We first examined the extent of changes in concrete indicators of SES, finding that there was little evidence for social mobility during the study period: maternal education did not change significantly over time ($F_{s(child\ age)}$ = 1.30, $p$ = 0.36), nor did income-to-needs ratio ($F_{s(child\ age)}$ = 1.01, $p$ = 0.49), nor did neighborhood deprivation (ADI, $F_{s(child\ age)}$ = 1.45, $p$ = 0.19). We next examined whether our results were robust to using a disadvantage factor score from later timepoints, rather than the prenatal disadvantage factor score.

We find that across timepoints, global segregation ($F_{s(agexSES\_y1)}$ = 2.72, $p$ = 0.07; $F_{s(agexSES\_y2)}$ = 3.32, $p$ = 0.013; $F_{s(agexSES\_y3)}$ = 6.785, $p$ = 0.0013), meso-scale segregation ($F_{s(agexSES\_y1)}$ = 4.19, $p$ = 0.016; $F_{s(agexSES\_y2)}$ = 5.87, $p$ = 0.003; $F_{s(agexSES\_y3)}$ = 7.56, $p$ = 0.0006), and local segregation (***Supplemental Figure 3d-f,*** $F_{s(agexSES\_y1)}$ = 4.83, $p$ = 0.005; $F_{s(agexSES\_y2)}$ = 8.71, $p$ = 0.0002; $F_{s(agexSES\_y3)}$ = 8.91, $p$ = 0.0002) all show similar patterns of interactions, such that infants and toddlers from lower-SES backgrounds show a faster increase in cortical network segregation than infants and toddlers from higher-SES backgrounds. We found some evidence for moderating effects of SES on developmental changes in network integration when using disadvantage factor scores from later timepoints ($F_{s(agexSES\_y1)}$ = 2.22, $p$ = 0.04; $F_{s(agexSES\_y2)}$ = 3.34, $p$ = 0.03; $F_{s(agexSES\_y3)}$ = 2.43, $p$ = 0.03).

The magnitude of SES effects on developmental increases in local segregation differed across functional systems, with the strongest effects found in somatomotor-hand, visual, somatomotor-mouth, dorsal attention, and parieto-occipital systems (***Supplemental Figures 4g-i***).

**Supplemental Table 1. Participant demographics at birth.**

| Variable | N = 261[1] |
|---|---|
| Age at scan (months) | 41.3 (38.0 - 45.0) |
| Child sex | |
| Male | 141 / 261 (54%) |
| Female | 120 / 261 (46%) |
| Gestational age (weeks) | 38.9 (37.0 - 41.6) |
| Birthweight (g) | 3,274.0 (2,200.0 - 4,627.0) |
| Area Deprivation Index | 67.5 (6.0 - 100.0) |
| Income to Needs Ratio | 2.8 (0.4 - 12.1) |
| Highest level of parent education completed | |
| Less than 12th grade | 22 / 251 (8.8%) |
| High school degree/GED | 101 / 251 (40%) |
| Some college/vocational school | 44 / 251 (18%) |
| College degree (4 years) | 29 / 251 (12%) |
| Graduate degree | 55 / 251 (22%) |
| Socioeconomic disadvantage factor score | -0.1 (-2.2 - 1.5) |

[1] Mean (Range); n / N (%)

### Supplemental Table 2. Bivariate correlations between SES variables at birth.

| | *Area Deprivation Index (ADI)* | *Income-to-Needs Ratio (INR)* | *Healthy Eating Index* | *Insurance status* | *Maternal education* |
|---|---|---|---|---|---|
| **Area Deprivation Index (ADI)** | **1.00** | | | | |
| **Income-to-Needs Ratio (INR)** | **-0.62*** ** | **1.00** | | | |
| **Healthy Eating Index** | **-0.28*** ** | **0.34*** ** | **1.00** | | |
| **Insurance status (private/individual or public/uninsured)** | **-0.53*** ** | **0.67*** ** | **0.27*** ** | **1.00** | |
| **Maternal education** | **-0.61*** ** | **0.71*** ** | **0.40*** ** | **0.59*** ** | **1.00** |

******* $p < 0.001$

Non-parametric statistic used for ordinal variables.
